# High-Low training is safe and effective in improving outcomes in a rodent model of chronic cervical spinal cord injury

**DOI:** 10.64898/2026.04.06.716770

**Authors:** D.R.S. Britsch, K.M. Cotter, C.M.J. Stuart, J. Turchan-Cholewo, M.K. Colson, E.D. Winford, T.A. Ujas, J. Lutshumba, A.L. Trout, C. Calulot, J.C. Gensel, W.J. Alilain, A.M. Stowe

## Abstract

Repeated exposure to hypoxia (oxygen levels below sea-level atmospheric conditions, ∼21%) alternated with regular voluntary exercise, known colloquially as ‘Living High, Training Low’, or simply ‘High-Low’, is used by elite athletes to boost exercise benefits and athletic performance. While paradigms of High-Low training have been utilized by Olympic athletes for decades, the therapeutic potential of a High-Low regimen in the context of neurotrauma has yet to be investigated. This long-term experiment evaluated the independent and combined effects of repeated hypoxic exposure and voluntary exercise on functional outcomes within the context of preclinical spinal cord injury (SCI). We hypothesized that combinatorial High-Low training enhances functional recovery, beyond either exercise or repeated exposures to hypoxia alone, to improve outcomes after SCI. Adult female rats (n=62) underwent a high-cervical hemisection (LC2H) to model spinal cord injury. At 6 weeks post-SCI, treatment (access to exercise wheel, repeated exposure to normobaric hypoxia at rest, or alternation of both) began in the surviving subjects (n=49). Despite initiation of treatment beyond the acute post-injury phase, High-Low therapy significantly improved respiratory function and prevented the development of SCI-associated anxiety-like behaviors. Notably, repeated *in vivo* exposure to normobaric hypoxia induced a shift in peripheral T cell profiles, characterized by increased CD4^+^ and reduced CD8^+^ expression. These findings indicate that combining repeated exposure to hypoxia with voluntary exercise as a therapy could promote recovery in the existing spinal cord-injured population. Collectively, this work provides a foundational first step for further investigation of High-Low training as a rehabilitation therapy for individuals living with SCI.

## INTRODUCTION

There are an estimated 300,000 people living with spinal cord injury (SCI) in the United States, with fewer than 1% achieving complete recovery at discharge.^1^ The recovery period following SCI occurs over months to years, but unfortunately, there are no FDA-approved treatments to reverse the damage caused by SCI.^2, 3^ Long-term effects of SCI are seen across multiple organ systems, including locomotor deficits, bowel and bladder dysfunction, respiratory dysfunction, cardiovascular complications, mental health decline, and chronic inflammation, most of which contribute to earlier mortality.^1–5^ The spinal cord maintains a degree of plasticity after traumatic injury, yet studies reporting robust improvement after intervention often focus on a single dysfunction despite the high level of interdependence between physiological systems. The lack of standardized care, coupled with the potential for recovery in the months to years following SCI, represents a gap in treatment for the heterogenous spinal cord-injured population.

Physical therapy is a core component of post-SCI rehabilitation, widely recognized for its dose-dependent benefits on functional recovery and overall health. Task-specific trainings can enhance performance in the measured outcome but may fail to support (or even limit) broader functional recovery in animal models.^6, 7^ In contrast, whole-body exercises engage multimodal plasticity and effectuate cortical reorganization to support global improvements.^8, 9^ Preclinical studies demonstrate that voluntary exercise initiated acutely after neurotrauma (∼2 weeks) improves locomotor outcomes, whereas overly intensive regimens (e.g., forced-use treadmill) or hyperacute initiation of exercise therapy (within the first 7 days) can exacerbate deficits.^6, 10–12^ In clinical populations, early initiation of rehabilitation is constrained by polytrauma, medical instability, and advanced age at injury as the average age of SCI patients continues to rise.^1, 13–15^ These factors underscore the need for adaptable exercise regimens that can be safely implemented after the acute phase across diverse patient populations.^16^ Although the mechanisms underlying physical therapy-induced recovery remain incompletely defined, exercise is known to upregulate neurotrophic growth factors, modulate inflammatory signaling, and promote CNS plasticity, even in individuals with chronic SCI.^17^ A growing body of evidence further suggests that augmenting exercise with other plasticity-enhancing interventions^18, 19^ or sublethal/stimulatory challenges^20^ can amplify its therapeutic efficacy.

The idea of combining exercise with exposure to hypoxia is not new; for decades, Olympic athletes have utilized “Living High, Training Low”, a training regimen involving acclimatization to high-altitude living coupled with exercise training at sea level. “Living <u>High</u>, Training <u>Low</u>”, or simply “High-Low”, increases exercise performance in elite athletes and is distinct from training *while* at high altitude or under hypoxic conditions, with recent studies suggesting that non-elite athletes may see even more pronounced benefits.^21–23^ However, temporary high-altitude living is not practical for non-elite athletes, or for the purposes of rehabilitation, hence treatment with exposure to hypoxia at sea-level has emerged as an alternative to replace “Living High”.^21–23^ A High-Low training regimen using voluntary exercise alternated with repeated hypoxia treatment represents a non-invasive, modifiable therapy with high translational potential to alleviate SCI-induced multi-system dysfunction in the existing pool of people living with SCI. We hypothesized that adapting High-Low training into a rehabilitative therapy would promote functional recovery compared to control subjects, beyond exercise or hypoxia treatment alone. Thus, we combined overnight access to running wheels with sustained hypoxic exposures administered during periods of rest to enhance recovery in a translational model of chronic-stage, cervical SCI.

## METHODS

### SCI – LC2H

Female Sprague-Dawley rats (Envigo-Harlan hsd:SD) acclimated to the housing facility for 1-2 weeks before undergoing assay training/baseline measures. Rats (age 3-4 months, n=62) received a left C2 hemisection (LC2H) using aseptic technique and a 3-pass method with a 27G syringe tip as previously described.^24^ Weight was recorded and anesthesia was induced with 4-5% isoflurane, after which depth of anesthesia was maintained at 2-3% isoflurane and monitored via toe-pinch reflex and observed breathing pattern. Buprenorphine SR (0.45mg/kg) was administered subcutaneously just prior to the end of anesthesia. Topical Bupivacaine (0.5mg/mL) was applied to the incision site immediately prior to incision and then again following staple closure. Once isoflurane anesthesia was halted, rats were placed in a clean cage with alpha-dri bedding on a water-circulation heating pad, followed by administration of subcutaneous saline and Carprofen (5mg/kg). Rats were monitored closely until fully awake and then given acute post-operative care twice daily for the first 3 days post-injury, or longer if necessary. Acute post-operative care includes administration of saline, daily weighing, supplemental nutrition (DietGel Recovery packs, 72-06-5022, Clear H_2_O, Westbrook, ME, USA), monitoring of surgical site, and additional doses of Buprenorphine (at 24hrs) and Carprofen (at 24, 48hrs). 12 subjects expired during the surgery or within the first 48 hours, with 50 surviving beyond the hyperacute/post-operative phase (survival rate of 81%). Staples were removed 10-14 days after surgery. After post-operative care was complete, weekly examinations and weigh-ins were recorded until the endpoint.

### Group Allocations

*A priori* exclusion criteria before group allocation included post-surgery weight loss greater than 20% (n=0), persisting bowel/bladder dysfunction (n=0), and paw autophagy (n=1). At 4 weeks post-injury (WPI), subjects (n=49) were assigned to groups using a random number generator. Our initial cohorts included only High-Low and Sedentary Control subjects. We incorporated crossed-control subjects (only one treatment) in the later cohorts; final counts were High-Low group n=16, Sedentary Control group n=16, Hypoxia-only group n=9, and Exercise-only group n=8. Standard Allentown IVC cages were used to pair-house subjects, except for one cage containing 3 rats in the Hypoxia-only group due to an uneven survival number for that surgical cohort.

### Hypoxia Administration (Living High)

Beginning at approximately 6WPI, subjects in High-Low or Hypoxia-only groups began a regimen of repetitive sustained hypoxia. Following return from overnight single-housing to pair-housing in the morning, High-Low or Hypoxia-only subject cages were sealed and had 11% O_2_ (custom blend oxygen tank 11% balanced with nitrogen, AWG Gases, Lexington, KY, USA) piped in at a rate of 1.5 L/hour for 4 hours, 5 days a week. Rats were housed in standard light/dark cycle so that the morning administration of hypoxia occurred during a period of rest/inactivity. Testing with an O_2_ sensor (Oxygen Detector, Forensics Detectors FD-600-HA-O2, Rolling Hills Estates, CA, USA) showed that oxygen levels reached 11% within an hour of initiation. Cages still provided *ad libitum* access to food and water. For the first 2 cohorts, we concurrently treated control group subjects in the same manner by piping in 21% O_2_ from compressed gas tanks. Review following the completion of cohorts 1 and 2 indicated that our method of repetitive sustained hypoxia treatment was well tolerated without unanticipated side effects to health (i.e., signs of increased stress), so we subsequently eliminated 21% O_2_ treatment for Sedentary Control and Exercise-only groups.

### Voluntary Exercise Regimen (Training Low)

Beginning at approximately 6WPI, subjects in the High-Low or Exercise-only groups were provided with access to running wheels overnight for 5 days a week via single housing in monitored wheel cages (Scurry Rat Running Wheel/Chamber, Lafayette Instruments 0859S, Lafayette, IN, USA). Hypoxia-only and Sedentary Control group subjects were simultaneously single-housed overnight in the standard cages for the same 5 nights a week. The following morning, rats returned to their pair-housed standard cages. Exercise cages still provided *ad libitum* access to food and water.

### Behavior and Motor Outcomes

Data were collected the week prior to injury and then at 5, 9, and 13 weeks after injury, corresponding to one week before treatment and then 4 and 8 weeks of treatment. For all behavioral measures, cages were habituated in the room for 30-60 minutes before data collection. We utilized an open field assay with laser tracking capabilities (SuperFlex Open Field apparatus and Fusion software, Omnitech Electronics, Inc, Columbus, OH, USA) to collect locomotor information over 2-minute trials. Open field data was analyzed for anxiety and curiosity behavior as well as general motor capabilities using the accompanying Fusion software; briefly, a grid was created and the time spent in the peripheral versus center regions, as well as rearing events, were recorded by infrared laser beams mounted to the plexiglass container. For time spent in the perimeter, raw data for post-treatment timepoints (9, 13WPI) were normalized to pre-treatment values (5WPI) to better visualize changes in anxiety-like behavior relative to treatment. We utilized Buxco Small Animal Whole Body Plethysmography Pro apparatus (WBP) with Finepointe software (Data Sciences International, New Brighton, MN, USA) to collect respiratory measures in awake/unanesthetized subjects over a 45-minute period. The first 15 minutes were considered additional habituation to the WBP chambers, followed by data collection for 30 minutes.

### Terminal Diaphragm Electromyography (EMG)

A subset of subjects comprising the initial cohorts (High-Low n=8, sedentary control n=8) underwent a terminal diaphragm EMG procedure to record bilateral muscle activity. Data was collected using the Spike2 software (Cambridge Electronic Design, Ltd, Cambridge, UK). Rats were anesthetized via 4% isoflurane induction followed by 2-3% continuous isoflurane through a nose cone, within faraday cages. A laparotomy exposed the diaphragm and 2 bipolar electrodes were placed laterally near the phrenic nerves to measure activity from the ipsi- and contralateral hemidiaphragm. Ground wires for each side were inserted between the skin and intercostal muscles. The abdomen was sutured and an acclimation period of 10 min elapsed from initial time of anesthesia before a 15-minute baseline measurement of diaphragmatic activity was recorded. A 15-second nasal occlusion was performed to elicit maximal output, followed by another 15 minutes of recorded activity.

### Perfusion and Sample Collection

Immediately following EMG completion, rats were euthanized via isoflurane overdose. Rats that did not receive a terminal EMG were also euthanized via isoflurane overdose. The abdominal cavity was opened and the splenic artery clamped before spleen was rapidly removed. Spleens were immediately placed in approximately 5mL complete media on ice. Blood was immediately collected into a 1.5mL Eppendorf tube containing 25uL heparin (100u/L) via puncture of the right atria. Transcardiac perfusion was subsequently performed by flushing 300mL ice-cold PBS into the aorta through a 22G syringe tip clamped in place with hemostat tool. Blood collected from cardiac puncture was left on ice for approximately 30-60 minutes after collection until centrifugation (1600*g*, 15 minutes, 4℃) and isolation of plasma. After perfusion, the following tissues were collected and flash-frozen using liquid nitrogen: brain, heart, lungs, soleus muscle, gastrocnemius muscle. Samples were stored at -80°C.

### Immune Cell Isolation and Flow Cytometry

Spleens were processed into a single cell suspension of leukocytes. Briefly, spleens were mechanically homogenized through a 70µm mesh filter with 15mL complete media (Supplemental Table 1). Homogenates underwent sequential centrifugation in media to wash and pellet cells (438*g*, 10 minutes, 4℃), and collection of lymphocytes was accomplished via density gradient using Lympholyte Rat Cell Separation Media (CedarLane CL5045, Burlington, NC, USA), ending in a single cell suspension. Samples from each cohort were banked in freezing media (Supplemental Table 2) and placed in a Mr. Frosty Freezing Container (ThermoFisher 5100-0001) to be stored at -80℃ for 48 hours, followed by transfer to liquid nitrogen storage at approximately -180℃ until unbanked.

Following completion of the final cohort samples were unbanked, tested for viability, and counted using Nexcelom ViaStain AO/PI (Revvity, CS2-0106-25mL) with Nexcelom Cellometer K2 (Revvity, 8003393 Rev E). One million cells per sample were aliquoted and stained with a panel of flow cytometry antibodies to characterize immune cell populations based on size, granularity, and marker expression (Supplemental Table 3). Using a custom BD FACSymphony A3 flow cytometer, 100,000-150,000 events per subject sample were recorded for analysis.

### Data acquisition and analysis

All data were collected using blinded subject numbers and then unblinded for analysis in GraphPad Prism (v9.1.0-10.6.1). Weekly weights were analyzed using 2way repeated measures (RM) ANOVA with Dunnett’s multiple comparisons correction, treatment groups vs. sedentary control group weeks 1-14. Exercise outcomes were analyzed using 2way RM ANOVA with Dunnett’s correction, week 1 vs week 2-8. Respiratory data were analyzed using 2way RM ANOVA with Dunnett’s correction, treatment groups vs. sedentary control group at 5, 9, 13WPI. Diaphragm EMG data were quantified using blinded subject numbers and raw traces in MATLAB and then unblinded for analysis in GraphPad Prism (v10.6.1) using 2way ANOVA with Sidak’s multiple comparisons correction, where EMG area under the curve (AUC) represents integrated, summative signal measured by the electrodes. Open field data were analyzed using 2way RM ANOVA with multiple comparisons and Dunnett’s correction, treatment groups vs. sedentary control group. Raw values of time spent in perimeter at 9, 13WPI were normalized to 5WPI values to better visualize changes in behavior pre-vs. post-treatment. Flow cytometry data was acquired by a single user with blinded subject numbers and single-color controls were simultaneously acquired to generate a compensation matrix within BDFACSDiva software (v8.0, BD Biosciences). FACS files and acquired compensation matrix were then exported and cell population gating was performed in FlowJo (v10.10, FlowJo LLC). Cell population data were analyzed using 2way RM ANOVA with multiple comparisons and Dunnett’s correction, treatment groups vs. sedentary control. See Supplemental Materials for full statistic sheets.

## RESULTS

### Rats with chronic, cervical SCI will exercise voluntarily

The experimental overview in Figure 1 depicts the timeline for spinal cord injury, treatment regimens and behavioral assays (Figure 1A) and a graphic of the LC2H including a representative section of the injury site stained with cresyl violet (Figure 1B). Weights were recorded during weekly health checks and the 4 groups in this experiment maintained similar post-SCI weight trajectory except at the week 14 endpoint, wherein the Exercise-only group became significantly heavier than the sedentary control group (Figure 1C). Our adaptation of High-Low training relied on voluntary exercise via overnight housing in wheel cages, which recorded distance and speed until the session was halted the following morning. The distances recorded each night were normalized to the length of each session due to overnight access to running wheels ranging from 12-15 hours. Normalized distances of the 5 sessions per week were then averaged for weekly representation. We saw no difference in the distance traveled between the High-Low and Exercise-only groups, but both groups did significantly increase average session distance over the course of 8 weeks (Figure 1D). Maximum speed was also recorded for each session and interpreted here as a measure of exercise intensity achieved. We again saw no difference between the High-Low and Exercise-only groups (Figure 1E), but a significant increase in maximum speed was attained by both groups over 8 weeks of wheel access.

**Figure 1:**
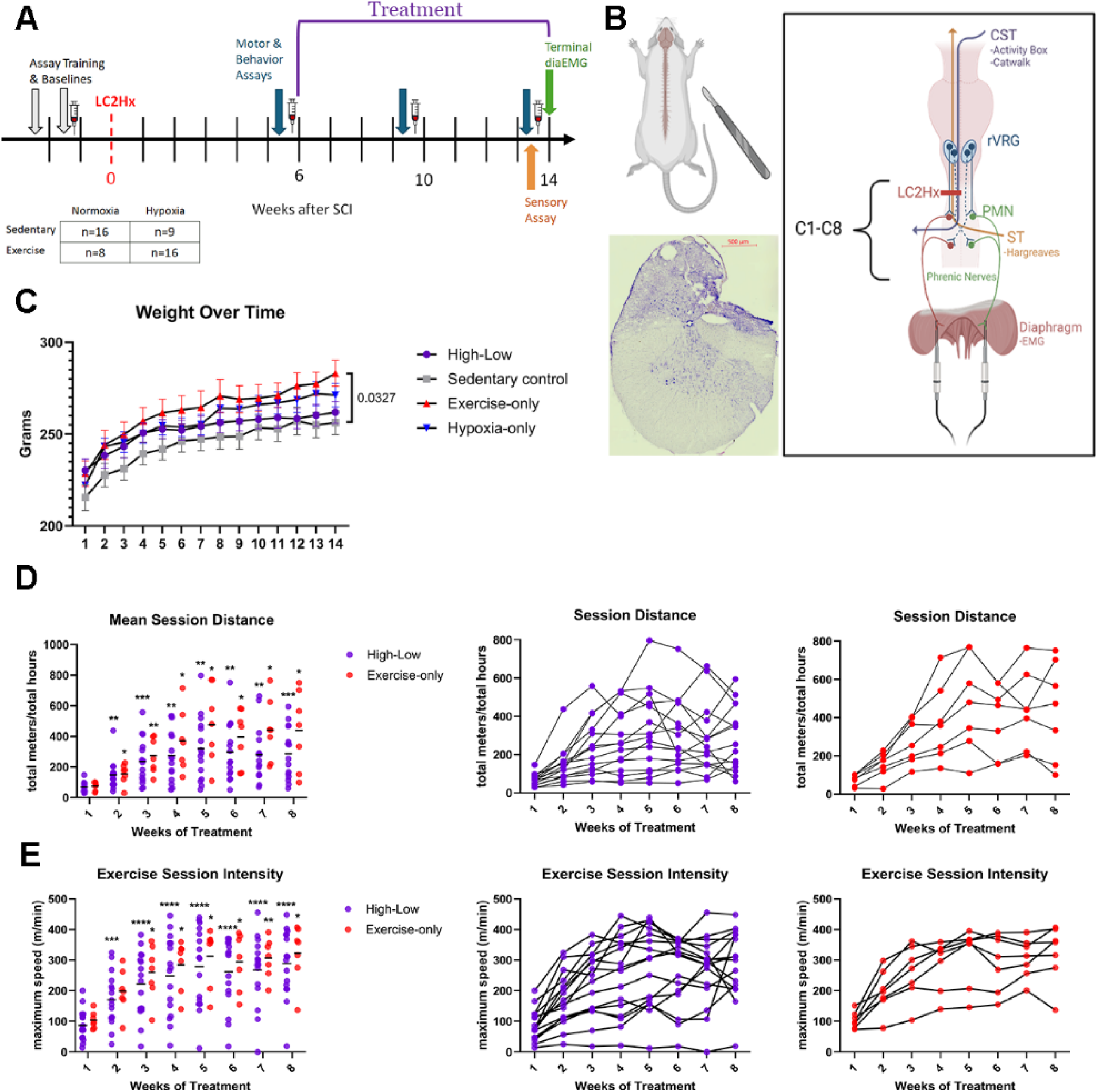
Experimental design, weekly weights and voluntary exercise output. Overview of experimental timeline (A). Diagram of left C2 hemisection model of spinal cord injury (generated with BioRender) and the resulting disruptions in relevant pathways, with representative image of injury stained with cresyl violet. Top to bottom; CST = corticospinal tract, rVRG = rostral ventral respiratory group, LC2Hx = left C2 hemisection, PMN = phrenic motor neurons, ST = spinothalamic tract, C1-C8 = cervical spinal levels 1-8 (B). Average weight ± SD (C). Distance traveled per session of wheel access, averaged across the week, with individual line plots show inter-subject variability (D). Maximum speed recorded within a given week, followed by individual line plots to show variability (E).

### High-Low and Hypoxia-only interventions alleviate post-SCI respiratory dysfunction

We utilized whole body plethysmography chambers to measure respiratory function in awake/unanesthetized rats. Most rats exhibited some degree of natural recovery by the endpoint, as evidenced by an increase in tidal volume and decrease in respiratory rate at 13WPI compared to 5WPI across all groups (Figure 2A-B). The recovery in tidal volume at 13WPI compared to 5WPI was significant in all groups except the Exercise-only group, which followed a similar trend but did not reach significance (Figure 2A). The High-Low group and Hypoxia-only group both experienced a robust increase in tidal volume after only 4 weeks of treatment, which persisted to endpoint (Figure 2A). Only the High-Low group demonstrated a corresponding significant decrease in respiratory rate after 4 weeks of intervention, although we saw a similar but non-significant trend in the Hypoxia- and Exercise-only groups (Figure 2B). A subset of High-Low and sedentary control cohorts ended with a terminal EMG recorded from the diaphragm (n=8/group). The low output of the ipsilateral hemidiaphragm, both AUC and peak inspiratory burst amplitude, indicates that hemisection injuries were functionally complete and little to no activity was recorded from the injured side during post-acclimation baseline measure (Fig 2C, 2D). Maximal output measured following a 15-second nasal occlusion indicated no significant difference in activity between baseline and occlusion measurements for either group, therefore failing to elicit activity on the injured side, although a higher degree of variability in response to nasal occlusion is apparent in the High-Low group (Figure 2E).

**Figure 2:**
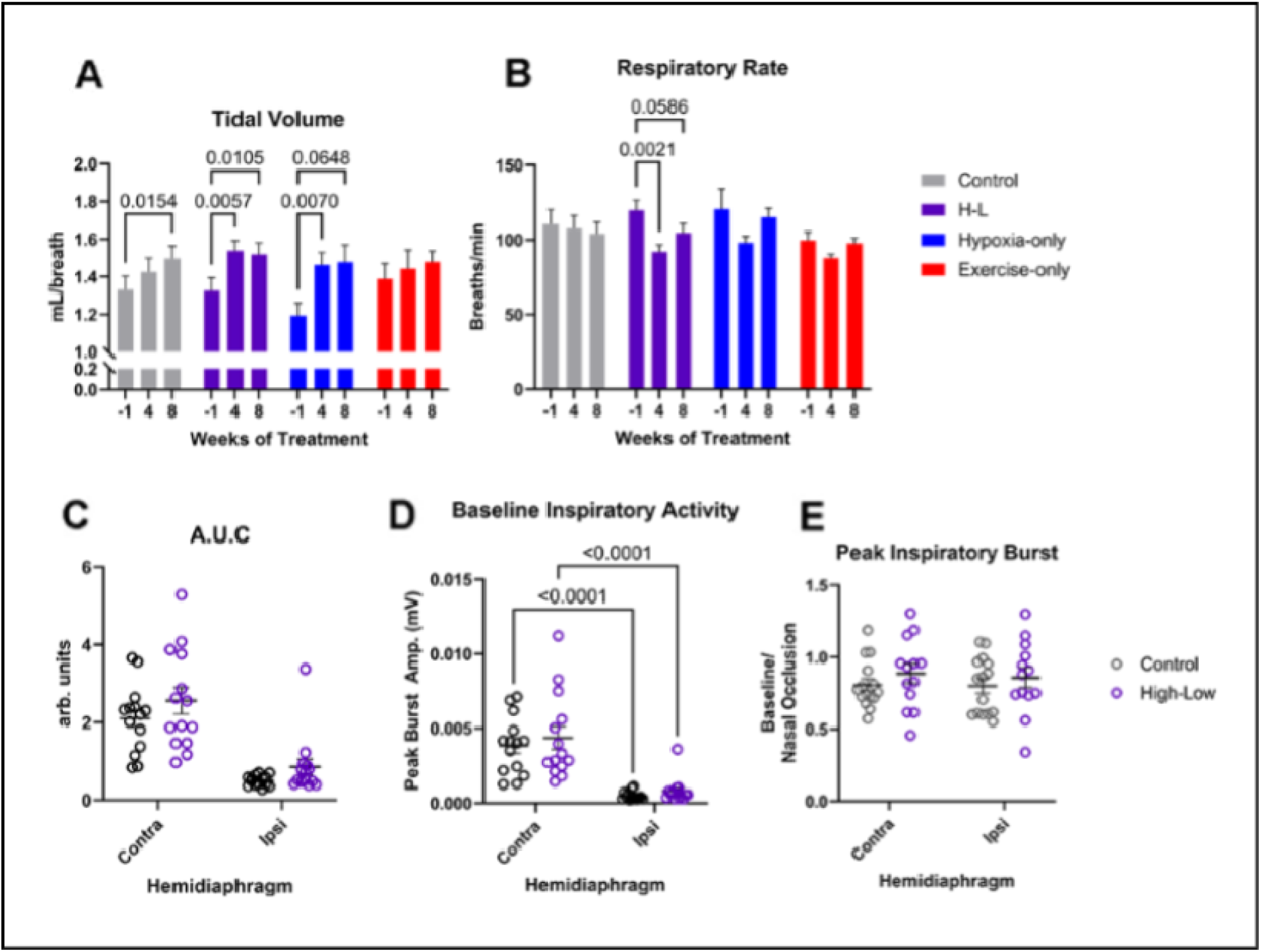
Hypoxia alleviates post-SCI respiratory dysfunction earlier than natural progression, but not through recovery of injured hemidiaphragm activity. Respiratory outcomes measured by whole body plethysmography ±SD (A,B) and terminal, bilateral diaphragm electromyogram ± SD (C-E).

### High-Low training eradicates anxiety-like behaviors post-SCI

The High-Low group exhibited reduced anxiety-like behavior at both 4- and 8-weeks of intervention, with these rats spending significantly less time in the perimeter than the Sedentary Controls (Figure 3A). Total distance traveled during the assay and number of rears were also recorded, providing additional data on exploratory behavior and gross locomotor ability. The High-Low group traveled significantly farther than the Sedentary Control group at 4 weeks of intervention (Figure 3B), while both High-Low and Exercise-only groups exhibited increased rearing compared to controls at the 4 week timepoint (Figure 3C).

**Figure 3:**
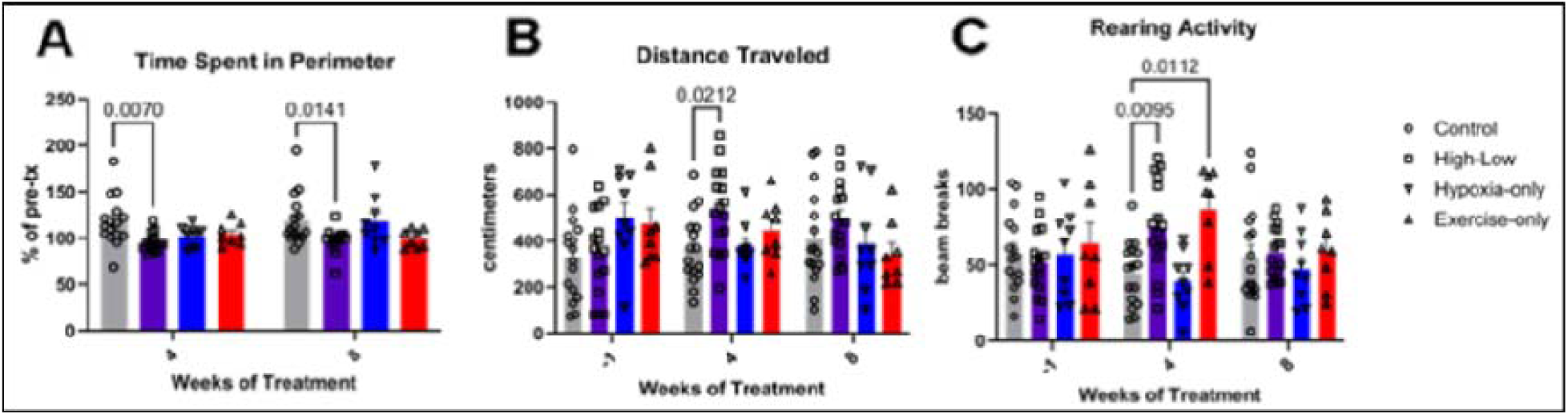
High-Low treatment eliminates anxiety-like behavior and increases exploratory behavior in the Open Field Assay. Time spent in perimeter of open field assay, normalized to pre-treatment timepoint as an indicator of anxiety-like behavior, ± SD (A).□Total distance traveled during open field assay, indicative of exploratory behavior and locomotive function, ± SD (B).□Number of rears recorded during open field assay, indicative of exploratory behavior and balance, ± SD (C).

### In vivo hypoxia alters splenic T cell populations

We isolated peripheral immune cells from the spleen at the 14WPI endpoint and subsequently performed flow cytometry. A custom panel (Supplemental Table 3) broadly characterized live leukocytes (Ghost Dye, granularity, and size) into lymphocytes (CD45), B or T lymphocytes (CD19 versus CD3), and T cell subtype (CD4 versus CD8; Figure 4A). Additional markers (granularity, size, His48, and CD11b) were used to identify myeloid-derived suppressor cells (MDSCs). While we did not see a difference in the overall proportion of B or T cells as a percentage of total CD45^+^ lymphocytes, or in MDSCs, a shift in T cell phenotypes was apparent (Figure 4B-D). In particular, the Hypoxia-only treatment increased the proportion of CD4^+^ helper cells while decreasing the proportion of CD8^+^ cytotoxic T cells (Figure 4E).

**Figure 4:**
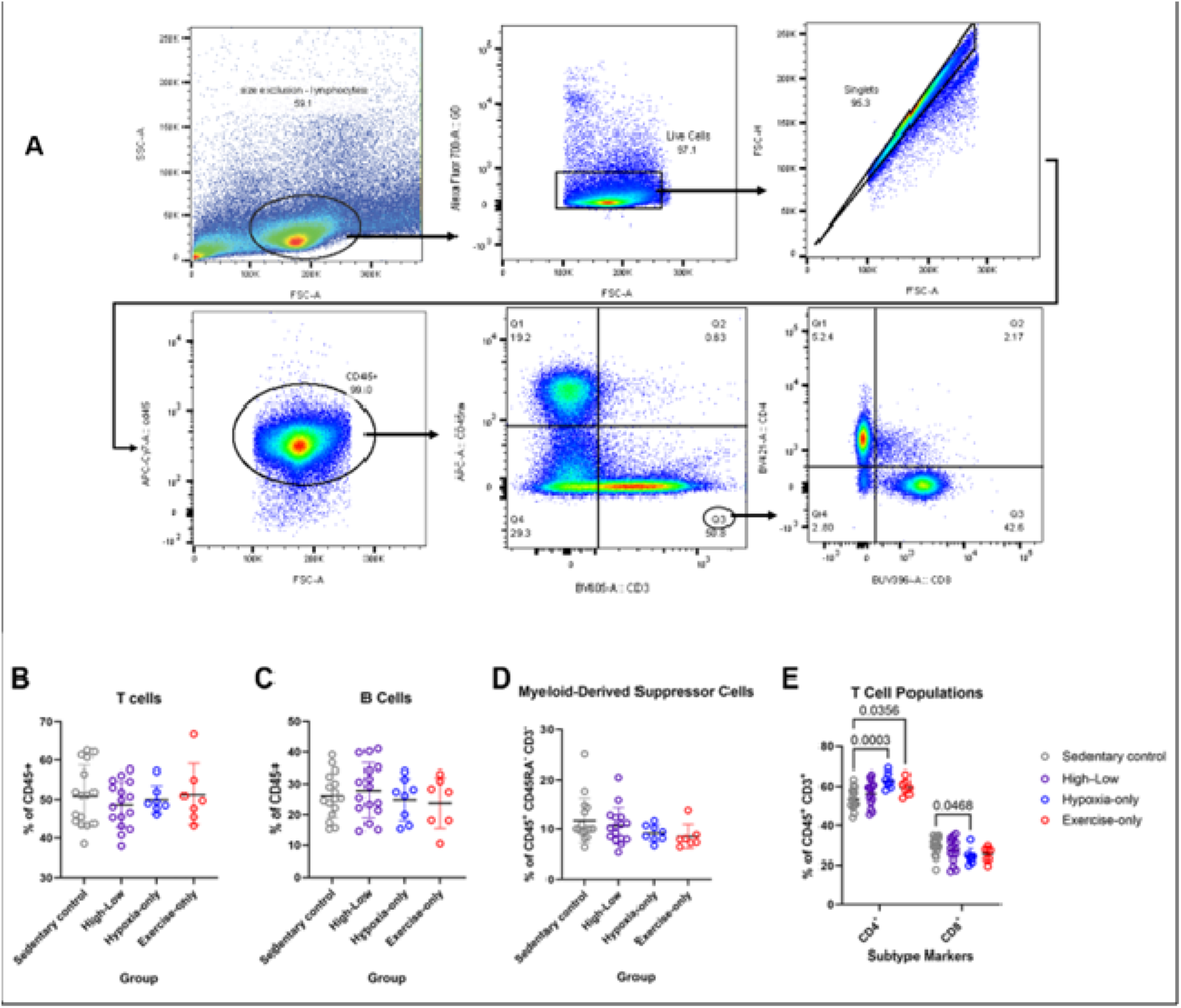
Characterization of immune cells via flow cytometry. Representative gating strategy to identify cell populations based on size, granularity, and negative or positive fluorescent compared to single-color controls with acquired compensation matrix (A). Proportion of T cells, B cells, and MDSCs ± SD (B-D). Proportion of CD4 versus CD8 positive T cells, mean plotted (E).

## DISCUSSION

Physical activity confers wide-ranging benefits for overall health and longevity, including in human SCI survivors, where exercise modulates inflammation and increases production of critical growth factors. Repetitive intermittent hypoxia treatment is one type of sublethal stressor which has been shown to improve respiratory outcomes following CNS injury by activating endogenous growth factor pathways and promoting neural plasticity.^25–30^ High-Low training, an existing regimen used by elite athletes, alternates exercise training with exposure to hypoxia while at rest to improve athletic performance. Here, we adapted a High-Low regimen for rats with chronic spinal cord injury, with a substantially delayed initiation of treatment post-SCI. Through the evaluation of our hypothesis that High-Low training would enhance recovery beyond either exercise therapy or hypoxia treatment alone, we observed specific outcomes that benefit from exercise or hypoxia treatment, and pleiotropic functional recovery in our High-Low group compared to untreated controls. Our findings highlight that repetitive, *sustained* hypoxia at rest can be used as a plasticity-enhancing treatment which synergizes with exercise rehabilitation to further promote recovery.

### Respiratory Outcomes

The LC2H injury model produces paralysis of the ipsilateral hemidiaphragm and respiratory impairment characterized by reduced tidal volume, increased respiratory rate, and impaired ventilation during hypercapnic challenge, with variable degrees of recovery occurring in approximately 50% of adult rats at 12WPI.^25, 31, 32^ This mirrors the prevalence and severity of respiratory dysfunction seen in humans, which is a leading cause of rehospitalization or death in SCI survivors.^1^ We therefore considered respiratory function our primary outcome and expected to see variable degrees of natural recovery across groups. Treatment with intermittent hypoxia can promote recovery of respiratory function after SCI,^25, 26, 29, 30^ but whether repeated, *sustained* bouts of hypoxia can elicit similar responses is unclear. In this study, we report that, following LC2H, average tidal volume recovered over time across all groups but was most pronounced in the High-Low and Hypoxia-only groups, particularly after 4 weeks of treatment. Importantly, only the High-Low group exhibited a significant reduction in respiratory rate, suggesting improved ventilatory efficiency. The Hypoxia-only group exhibited a similar trend, but smaller sample size may have limited statistical power. Together, these findings suggest that delayed, daily sustained hypoxia therapy—distinct from intermittent hypoxia—can also improve respiratory parameters in chronic SCI.

Bilateral EMG recordings did not demonstrate notable recovery of ipsilateral hemidiaphragm activity in the High-Low (or sedentary control) group. Others have reported similar observations of non-dysfunctional respiration via whole body plethysmography despite continued paralysis of the injured hemidiaphragm in the same model of SCI but at an acute timepoint.^33^ Compensatory mechanisms such as strengthening of preserved contralateral circuitry or increased participation of accessory breathing musculature could be contributing to improvements in respiratory measures via whole body plethysmography without restoration of ipsilateral diaphragm function. Future studies incorporating bilateral recordings of the intercostal muscles, assessment of phrenic nerve activity, and quantification of serotonergic fibers will be essential to delineate the anatomical substrates of respiratory recovery following High-Low therapy.

### Voluntary Exercise and Metabolic Effects

Although thoracic and lumbar injury models are more commonly used to assess locomotor rehabilitation in preclinical work, cervical SCI remains the most common and clinically devasting level of injury,^1^ and is therefore important to investigate. To our surprise, all injured subjects exercised voluntarily (albeit to various degrees) despite high cervical injury and chronic forelimb impairments. The variability we saw in initial capacity, voluntary continued use, and improvement in exercise outcomes is highly representative of the variability in human exercise inclination and capacity.^34^ Both High-Low and exercise-only groups demonstrated progressive increases in running distance and maximum speed across the 8-week rehabilitation period, with no significant difference between the two groups. We can therefore conclude that daily sustained hypoxia at rest did not adversely affect voluntary exercise performance. An unexpected finding was significantly greater terminal body weight in the Exercise-only group, but not the High-Low group, compared to the Sedentary Control group. Weight gain over time is expected in Sprague-Dawley females,^35^ and the propensity for exercise to minimize their weight gain is mitigated by compensatory eating,^36^ which may explain why the Exercise-only group averaged the heaviest weight despite regular activity. The fact that terminal weight was not significantly higher in the High-Low group suggests a potential metabolic interaction between exercise and sustained hypoxia. This aligns with recent clinical evidence that sustained overnight hypoxia aids weight loss by increasing resting metabolic rate and reducing appetite,^37^ with the important distinction that the human subjects had a BMI of 30 or greater at the start of the study. Future investigations should verify similar metabolic effects in non-obese rodents exposed to sustained hypoxia (with or without exercise) using monitored feeding and/or metabolic cages.

### Affective and Exploratory Behavior

Neuropsychiatric symptoms, particularly anxiety and depression, frequently emerge after cervical SCI in both clinical and experimental settings.^38–41^ Here, we measured affective behavior in post-SCI rats utilizing a classical open field assay with added laser-tracking to collect additional rearing and movement data. We observed a progressive increase in anxiety-like behavior in the Sedentary Control group over the course of 8 weeks. In contrast, the High-Low group traveled farther distances and exhibited increased rearing at 9WPI, spending less time in the perimeter at both post-treatment timepoints (9, 13WPI) compared to the pre-treatment baseline. These findings suggest that our novel adaptation of delayed High-Low training as a rehabilitation therapy prevented the development of SCI-associated anxiety behavior.

Exercise-only treatment also prevented the injury-associated increase in anxiety-like behavior but did not demonstrate the same degree of reduction as the High-Low group. Like the High-Low group, the Exercise-only group performed more rears than the Sedentary Control group at 9WPI but did not travel significantly farther. We postulate that exercise is the driving factor behind improvements in the current study but there is a clear synergistic effect seen in the High-Low group. As this assay simultaneously measures multiple parameters, it is important to note that we cannot explicitly distinguish to what extent the observed reduction of anxiety-like behavior is due to improvement in cognitive state versus locomotor capabilities. A salient study investigating anxiety and depression in spinally contused rats illuminates the complex relationship between anxiety, depression and locomotion after injury, highlighting the fact that the two negative affective behaviors have distinct inflammatory profiles yet are not intrinsically linked with severity of injury.^42^ This supports our emphasis of High-Low training’s ablation of increased anxiety-like behavior post-SCI, regardless of potential differences in locomotive ability.

### Adaptive Immunomodulation

B and T lymphocytes interact and coordinate with one another to mount an adaptive immune response, which can last for months to years after neurotrauma.^43–45^ Infiltration of T cells into the lesion after SCI has been observed via immunohistochemistry across several animal models.^46^ While the exact role of T cells in the lesion remains controversial, CD8^+^ T cells are recruited to the injury site through the CCL2/CCR2 chemokine axis and have been shown to exacerbate secondary injury through production of perforin.^46^ A prospective clinical study confirmed that CD8^+^ T cell populations are increased in the blood of SCI patients over the first 3 days post-injury.^45^

Preclinical studies have reported highly variable effects of physical activity on T cell populations, dependent on species, age, source of cells, and intensity/duration of exercise regimen. One such study in adult Wistar male rats showed that 4 weeks of moderate exercise increased splenic CD8^+^ T cell populations but no change in CD4^+^ was observed,^47^ while another study in male and female C57Bl/6 mice showed that voluntary wheel running increased lymph node-derived CD4^+^ T cells in young, but not adult or old, subjects.^48^ We have previously reported that high-volume exercise is inversely correlated to CD4^+^ *and* CD8^+^ T cells in the naïve mouse brain, while no differences to splenic populations were observed in the exercise versus sedentary groups.^49^

Immune cells within niches such as the bone marrow and intestines are known to be regulated by hypoxia (primarily through HIF signaling) under normal physiological conditions, but the effect of global hypoxia on peripheral immune cells is less characterized, especially in the context of spinal cord injury.^50, 51^ We previously reported that repeated, sustained exposure to hypoxia as a preconditioning stimulus before ischemic stroke in mice induces an immunosuppressed phenotype in B cells and reduces the quantity of CD4^+^ T cells within the post-stroke brain.^52^ To determine whether our adaptation of High-Low training alters circulating immune cell populations in post-SCI rats, we isolated peripheral lymphocyte populations and characterized them via flow cytometry.

Our analysis of immune cells from the spleens of adult, female rats at 14WPI identified a shift in T cell populations in the single-treatment groups. We did not see changes in the overall proportion of B or T cells within the general lymphocyte population, nor did we see differences in MDSCs, which are immature myeloid cells that significantly expand during inflammation and strongly regulate T cell responses (Figure 4B-D).^53^ MDSCs are an innate immune cell that participate in rapid immune response and we did not expect to see differences in splenic populations nearly 4 months after the injury occurred. Exercise-only treatment increased the proportion of CD4^+^ T cells while CD8^+^ T cells appear unaffected (Figure 4E). The most intriguing observation was that Hypoxia-only treatment increased CD4^+^ helper T cell representation *and* decreased CD8^+^ cytotoxic T cell representation (Figure 4E). To our knowledge, this is the first demonstration of hypoxia-induced modulation of systemic T cells in a rat model of SCI, and we find this result particularly interesting considering human SCI patients have exhibited decreased CD4^+^ helper T cells and increased CD8^+^ cytotoxic T cells acutely after injury.^45^ This observed shift supports the concept that non-invasive hypoxia therapy may exert long-term immunomodulatory effects in chronic SCI. Planned immunohistochemistry will investigate immune cells near the injury site. Future follow-up studies will further characterize and evaluate hypoxia-conditioned lymphocytic responses to inflammatory challenges.

## CONCLUSIONS

While combinatorial therapeutic strategies are not inherently novel, this study is, to our knowledge, the first to implement a High–Low training paradigm in a preclinical model of spinal cord injury. We hypothesized that using repetitive, sustained hypoxia as a plasticity-enhancing, sublethal stressor alongside exercise rehabilitation (i.e., High-Low) would synergistically promote recovery. This interaction was most evident in respiratory function and anxiety-like behavior, where the combined intervention outperformed either modality alone. In other domains, exercise and hypoxia appeared to exert differential effects, consistent with prior reports of their independent benefits.^17, 26, 54–56^ Importantly, this regimen was initiated 6 weeks after injury, beyond the acute and subacute phases often targeted in experimental SCI studies investigating exercise therapy. Despite this delay, High–Low training was safe, well-tolerated, and associated with improvements spanning respiratory physiology, exploratory behavior, and adaptive immune composition. The observed shift in splenic T cell populations following sustained hypoxia further suggests that systemic physiological interventions can reshape immune profiles in chronic SCI, an area that remains incompletely defined in translational neurotrauma research.

Collectively, these findings support the premise that the chronically injured spinal cord remains biologically dynamic and responsive to non-invasive, combinatorial rehabilitation strategies. The voluntary nature of the exercise component enhances translational feasibility, allowing individualization based on initial capacity and progressive improvement. By targeting multiple systems simultaneously and extending the therapeutic window into the chronic stage, High–Low training represents a promising framework for rehabilitation in the existing population of SCI survivors. Thus, the main conclusions from this study are (1) that High-Low training is a safe, well-tolerated regimen after SCI in rodents, and (2) effectively promotes holistic recovery beyond the acute post-injury stage.

### TRANSPARENCY, RIGOR, AND REPRODUCIBILITY

*Post-hoc* power analysis of our *a priori* primary outcome, respiratory measures, confirmed for the continuous endpoint two independent sample study (High-Low versus Sedentary control, n=16/group) a 99.2% power achieved with an α=0.05. Efforts to maintain rigor were incorporated at each stage of this work. Behavioral assays were performed under blinded conditions, with each subject receiving a number based on surgical cohort (i.e., cohort 1 subject 2) rather than a descriptor of treatment group. Control groups were used and included no treatment (i.e. sedentary control mice), as well as single-treatment (i.e., Exercise-only and Hypoxia-only). Pair-housed cages were randomly assigned to groups immediately prior to pre-treatment data collection timepoint (4-5WPI). Data were recorded using only the surgical cohort/subject numbers. Execution of assays included pre-training/baseline measures, appropriate habituation and acclimatization to rooms and equipment, and consistency of personnel performing assays. Whole body plethysmography, activity box and exercise output data were recorded automatically by each respective system and data were stored on LabArchives. After the end of cohorts, the subject number key was used to unblind groups for analysis in GraphPad Prism. A data analysis plan was not formally pre-registered but was pre-specified in the grant funding. This manuscript is available to the public via BioRxiv and raw data will be available on the SCI Open Data Commons website (see statements and declarations below). Lesion quantification is ongoing and will be added to the dataset upon completion.

## ACKNOWLEDGEMENTS

Acknowledgements and gratitude to Dr. Michael Sunshine for assisting with quantification of Spike2 EMG traces using his MATLAB code. Please note that since completion of this work, and at time of manuscript submission, the following co-authors have moved to different institutions:

Connor Stuart: Department of Neurology, University Medical Center Hamburg-Eppendorf, Hamburg, Germany

Dr. Winford: Department of Neurology, Taub Institute for Research on Alzheimer’s Disease and the Aging Brain, Gertrude H. Sergievsky Center, Vagelos College of Physicians and Surgeons, Columbia University, New York, NY, USA

Dr. Trout: Department of Neurology, University of Pittsburgh, Pittsburgh, PA, USA

## AUTHORSHIP CONTRIBUTIONS

DRSB: SCI surgeries, data acquisition, tissue collection, data analysis, manuscript drafting and editing; KMC.: data acquisition, tissue collection, manuscript edits; CJS: data acquisition, tissue collection, data analysis; JTC: data acquisition, tissue collection, manuscript edits; MKC: tissue collection; EDW: tissue collection, manuscript edits; TAU: tissue collection, manuscript edits; JL: tissue collection, manuscript edits; ALT: tissue collection, manuscript edits; CC: SCI surgeries; JCG: study conception and design, manuscript edits; WJA: study conception and design, SCI surgeries, manuscript edits; AMS: study conception and design, tissue collection, data analysis, manuscript edits.

## STATEMENTS AND DECLARATIONS

## Ethical Considerations

All surgeries, procedures, assays and housing were done in accordance with approved IACUC protocol (2018-3125), following the NIH Guide for the Care and Use of Laboratory Animals and with the support of the University of Kentucky Division of Laboratory Animal Resources.

## Consent to participate

not applicable

## Consent for publication

not applicable

## Data availability

This manuscript is available on BioRXIV beginning April 2026. We will upload raw datasets on ODC-SCI within 30 days of peer-reviewed publication.

Uncategorized References

