## Supplemental Tables for "High-Low training is safe and effective in improving outcomes in a rodent model of chronic cervical spinal cord injury"

**Table 1:** Recipe for RPMI-based complete media for single-cell suspension.

| **Reagent** | **Volume** | **Vendor** | **Catalog #** |
| --- | --- | --- | --- |
| RPMI 1640 (w/out L-glut) | 500mL | Corning | 10-040-CV |
| Fetal Bovine Serum (heat-inactivated) | 50mL | Sigma Aldrich | F4135-500 |
| 1M HEPES Buffer | 6.25mL | Gibco (ThermoFisher) | 15630080 |
| 10nM non-essential Amino Acids | 5mL | Corning | 25-025-CI |
| HyClone Penicillin-Streptomycin | 5mL | Cytiva | SV30010 |
| 100X GlutaMAX | 5mL | Gibco (ThermoFisher) | 35050061 |
| 100mM Sodium Pyruvate | 5mL | Gibco (ThermoFisher) | 11360070 |
| 14.3M 2-Mercaptoethanol | 1µL | Sigma | M3148-25 |

**Table 2:** Recipe for freezing media for cell storage in liquid nitrogen.

| **Reagent** | **Volume** | **Vendor** | **Catalog #** |
| --- | --- | --- | --- |
| Fetal Bovine Serum (heat-inactivated) | 250mL | Sigma Aldrich | F4135-500 |
| Complete media | 200mL | (Table 1) | n/a |
| DMSO | 50mL | Fisher | BP231-1 |

**Table 3:** Flow cytometry antibodies to characterize splenic immune cell populations.

| **Antibody** | **Fluorophore** | **Clone** | **Vendor** | **Catalog #** | **Lot #** |
| --- | --- | --- | --- | --- | --- |
| CD45 | APC-Cy7 | OX1 | BioLegend | 202216 | B373471 |
| CD45RA | APC | OX33 | BioLegend | 202314 | B387620 |
| CD3 | BV605 | 1F4 | BD | 563949 | 3180461 |
| CD4 | V450 | OX35 | BD | 561579 | 373107 |
| CD8a | BUV395 | OX-8 | BD | 740257 | 3255773 |
| HIS48 | FITC | HIS48 | Invitrogen | 11-0570-82 | 2685491 |
| CD11b/c | BV510 | OX42 | BD | 743878 | 325576 |
| Ghost dye | Red 710 | n/a | Tonbo Biosci | 13-0871-T100 | D087105232313 |

**Table 4:** Two-way RM ANOVA statistical interactions for exercise data (Fig. 1).

| **Weekly Weight (Fig. 1C)** | | | | |
| --- | --- | --- | --- | --- |
| **Source of variation** | **F (DFn, DFd)** | **% total variation** | **P value** |  |
| Time x Group | F (16.14, 242.2) = 2.712 | 0.8862 | 0.0005 | *** |
| Time | F (5.381, 242.2) = 175.9 | 19.17 | <0.0001 | **** |
| Group | F (3, 45) = 1.477 | 6.706 | 0.2336 | ns |
| Subject | F (45, 585) = 180.6 | 68.11 | <0.0001 | **** |
| **Mean Session Distance (Fig. 1D)** | | | | |
| **Source of variation** | **F (DFn, DFd)** | **% total variation** | **P value** |  |
| Time x Group | F (6, 88) = 1.220 | 2.222 | 0.0523 | ns |
| Time | F (1.584, 69.68) = 6.704 | 27.47 | <0.0001 | **** |
| Group | F (3, 44) = 0.9947 | 4.713 | 0.1654 | ns |
| Subject | F (44, 88) = 2.859 | 45.47 | <0.0001 | **** |
| **Exercise Session Intensity (Fig. 1E)** | | | | |
| **Source of variation** | **F (DFn, DFd)** | **% total variation** | **P value** |  |
| Time x Group | F (3.691, 77.52) = 0.1006 | 0.06898 | 0.9769 | ns |
| Time | F (3.691, 77.52) = 40.07 | 27.46 | <0.0001 | **** |
| Group | F (1, 21) = 0.6424 | 1.609 | 0.4318 | ns |
| Subject | F (21, 147) = 25.58 | 52.61 | <0.0001 | **** |

**Table 5:** Two-way RM ANOVA statistical interactions for respiratory data (Fig. 2).

| **WBP Tidal Volume (Fig. 2A)** | | | | |
| --- | --- | --- | --- | --- |
| **Source of variation** | **F (DFn, DFd)** | **% total variation** | **P value** |  |
| Time x Group | F (5.967, 89.51) = 0.9416 | 1.775 | P=0.4692 | ns |
| Time | F (1.989, 89.51) = 15.16 | 9.526 | P<0.0001 | **** |
| Group | F (3, 45) = 0.3138 | 1.211 | P=0.8153 | ns |
| Subject | F (45, 90) = 4.095 | 57.90 | P<0.0001 | **** |
| **WBP Tidal Volume (Fig. 2B)** | | | | |
| **Source of variation** | **F (DFn, DFd)** | **% total variation** | **P value** |  |
| Time x Group | F (6, 88) = 1.220 | 2.994 | 0.3039 | ns |
| Time | F (1.584, 69.68) = 6.704 | 5.486 | 0.0043 | ** |
| Group | F (3, 44) = 0.9947 | 3.491 | 0.4042 | ns |
| Subject | F (44, 88) = 2.859 | 51.47 | <0.0001 | **** |
| **EMG AUC (Fig. 2C)** | | | | |
| **Source of variation** | **F (DFn, DFd)** | **% total variation** | **P value** |  |
| Hemidiaphragm x Group | F (1, 26) = 0.04088 | 0.04074 | 0.8413 | ns |
| Hemidiaphragm | F (1, 26) = 48.44 | 48.27 | <0.0001 | **** |
| Group | F (1, 26) = 3.186 | 2.814 | 0.0860 | ns |
| Subject | F (26, 26) = 0.8866 | 22.97 | 0.6194 | ns |
| **EMG Mean Inspiratory Activity (Fig. 2D)** | | | | |
| **Source of variation** | **F (DFn, DFd)** | **% total variation** | **P value** |  |
| Hemidiaphragm x Group | F (1, 26) = 0.03673 | 0.03646 | 0.8495 | ns |
| Hemidiaphragm | F (1, 26) = 50.53 | 50.17 | <0.0001 | **** |
| Group | F (1, 26) = 0.7943 | 0.7110 | 0.3810 | ns |
| Subject | F (26, 26) = 0.9017 | 23.28 | 0.6030 | ns |
| **EMG Maximal Inspiratory Activity (Fig. 2E)** | | | | |
| **Source of variation** | **F (DFn, DFd)** | **% total variation** | **P value** |  |
| Hemidiaphragm x Group | F (1, 26) = 0.04335 | 0.07048 | 0.8367 | ns |
| Hemidiaphragm | F (1, 26) = 0.09947 | 0.1617 | 0.7550 | ns |
| Group | F (1, 26) = 1.254 | 2.647 | 0.2730 | ns |
| Subject | F (26, 26) = 1.298 | 54.87 | 0.2552 | ns |

**Table 6:** Two-way RM ANOVA statistical interactions for behavioral data (Fig. 3).

| **Anxiety Behavior (Fig. 3A)** | | | | |
| --- | --- | --- | --- | --- |
| **Source of variation** | **F (DFn, DFd)** | **% total variation** | **P value** |  |
| Time x Group | F (3, 45) = 3.069 | 2.344 | 0.1107 | ns |
| Time | F (1, 45) = 2.648 | 0.6742 | 0.0093 | ** |
| Group | F (3, 45) = 4.312 | 19.11 | <0.0001 | **** |
| Subject | F (45, 45) = 5.803 | 66.49 | 0.0373 | * |
| **Distance Traveled (Fig. 3B)** | | | | |
| **Source of variation** | **F (DFn, DFd)** | **% total variation** | **P value** |  |
| Time x Group | F (3, 45) = 0.6019 | 0.4477 | 0.6171 | ns |
| Time | F (1, 45) = 0.5758 | 0.1428 | 0.4519 | ns |
| Group | F (3, 45) = 3.702 | 17.49 | 0.0183 | * |
| Subject | F (45, 45) = 6.351 | 70.86 | <0.0001 | **** |
| **Rearing (Fig. 3C)** | | | | |
| **Source of variation** | **F (DFn, DFd)** | **% total variation** | **P value** |  |
| Time x Group | F (6, 90) = 4.792 | 9.316 | 0.0003 | *** |
| Time | F (1.959, 88.14) = 1.187 | 0.7695 | 0.3092 | ns |
| Group | F (3, 45) = 2.009 | 7.203 | 0.1261 | ns |
| Subject | F (45, 90) = 3.688 | 53.77 | <0.0001 | **** |

**Table 7:** ANOVA statistical interactions for immune cell data (Fig. 4).

| **MDSC’s** (Ordinary One-way) **(Fig. 4B)** | | | | |
| --- | --- | --- | --- | --- |
| **ANOVA table** | **F** | **R^2^** | **P value** |  |
| Groups | 1.670 | 0.1044 | P=0.1876 | ns |
| **CD45RA^+^ B Cells** (Ordinary One-way) **(Fig. 4C)** | | | |  |
| **ANOVA table** | **F** | **R^2^** | **P value** |  |
| Groups | 0.5186 | **0**.03492 | P=0.6718 | ns |
| **CD3^+^ T Cells** (Ordinary one-way) **(Fig. 4D)** | | | | |
| **ANOVA table** | **F** | **R^2^** | **P value** |  |
| Groups | 0.3857 | 0.02681 | 0.7639 | ns |
| **CD4^+^ vs CD8^+^ T Cells** (Two-way RM ANOVA) **(Fig. 4E)** | | | |  |
| **Source of variation** | **F (DFn, DFd)** | **% total variation** | **P value** |  |
| Cell type x Group | F (3, 43) = 5.061 | 2.728 | 0.0043 | ** |
| Cell type | F (1, 43) = 472.9 | 84.99 | <0.0001 | **** |
| Group | F (3, 43) = 1.389 | 0.1763 | 0.2589 | ns |
| Subject | F (43, 43) = 0.2354 | 1.819 | >0.9999 | ns |
